# *Fasciola hepatica* GST mu-class suppresses the cytokine storm induced by *E. coli*-lipopolysaccharide whereas modulates the dynamic of peritoneal macrophages in a mouse model and suppresses the classical activation of macrophages

**DOI:** 10.1101/2023.08.10.552847

**Authors:** Bianca N. Valdes-Fernandez, Caleb Ruiz-Jimenez, Albersy Armina-Rodriguez, Loyda B Mendez, Ana M. Espino

## Abstract

The helminth *Fasciola hepatica* is known as a master of immunomodulation. It suppresses antigen specific Th1 responses in concurrent bacterial infections while promoting the Th2/Treg regulatory responses, thus demonstrating its anti-inflammatory ability *in vivo*. We have recently demonstrated that a single intraperitoneal injection with native *F. hepatica* Glutathione *S*-Transferase (nFhGST), mostly comprised of mu-class isoforms, can suppresses the cytokine storm and increasing the survival rate in a mouse model of septic shock (1). Knowing that the peritoneal macrophages in response to microbial stimuli play essential roles in the defense, tissue repairment, and maintenance of homeostasis, the present study aimed to determine whether nFhGST could modulate the amount and dynamic of these cells concurrently to the suppression of pro-inflammatory cytokines. The remarkable findings described in this article are, (i) nFhGST suppresses serum IL-12, TNF-α, and IFN-γ in BALB/c mice challenged with a lethal dose of LPS, (ii) Although nFhGST does not elicit IL-10, it was able to significantly suppress the high levels of LPS-induced IL-10, which is considered a key cytokine in the pathophysiology of sepsis (2). iii) nFhGST prevent the disappearance of large peritoneal macrophages (LPM) whereas significantly increasing this population in the peritoneal cavity (PerC) of LPS treated animals, (iv) nFhGST promotes the alternative activation of macrophages whereas suppress the classical activation of macrophages *in vitro* by expressing high levels of Ym-1, a typical M2-type marker, secreting the production of IL-37, and preventing the production of TNF-α, iNOS2 and nitric oxide, which are typical markers of M1-type macrophages, (v) nFhGST suppress the bacterial phagocytosis of macrophages, a role that plays both, M1-and M2-macrophages, thus partially affecting the capacity of macrophages in destroying microbial pathogens. These findings present the first evidence that nFhGST is an excellent modulator of the PerC content *in vivo,* reinforcing the capacity of nFhGST as an anti-inflammatory drug against sepsis in animal models.

**Importance:** Sepsis is an infection that can lead to a life-threatening complication. Sepsis is the consequence of a systemic bacterial infection that exacerbates the immune cells’ activation by bacterial products, resulting in the augmented release of inflammatory mediators. A critical factor in the pathogenesis of sepsis is the primary component of the outer membrane of Gram-negative bacteria known as lipopolysaccharide (LPS), which is sensed by toll-like receptor 4 (TLR4). For this reason, scientists aimed to develop antagonists able to block the cytokine storm by blocking TLR4. We report here that a mixture of mu-class isoforms from the *F. hepatica* glutathione S-transferase (nFhGST) protein family administered intraperitoneally 1 h after a lethal LPS injection, is capable of significantly suppressing the LPS-induced cytokine storm in a mouse model of septic shock whereas modulate the dynamic and abundance of large peritoneal macrophages in the peritoneal cavity of septic mice. These results suggest that nFhGST is a prominent candidate for drug development against endotoxemia and other inflammatory diseases.

## Introduction

*Fasciola hepatica* one of the most prevalent parasitic helminths worldwide, is a master of immunomodulation. It is well documented that *F. hepatica* suppresses Th1 responses to establish long-lasting chronic infections in their mammalian hosts (3, 4). To achieve this effect from the earlier phase of infection, *F. hepatica* secrete a myriad of molecules called excretory-secretory products (ESPs), which suppress the Th1-immune response whereas concurrently promote the Th2/Treg response (3, 5, 6). *F. hepatica* infection has also been shown to attenuate the clinical symptoms of murine autoimmune encephalomyelitis (7) and prevent type-1 diabetes development in the non-obese diabetic mouse model (8). However, while live infection with *F. hepatica* is intensely immunoregulatory, it is impractical as a therapeutic option because these infections lack specificity and result in a compromised immune system unable to respond satisfactorily to bystander’s infections that require Th1 response for immunity (9, 10). Therefore, there is an urgent need to identify and characterize the immune-modulatory molecules produced by the parasite to develop feasible therapeutic options against inflammatory medical conditions.

Glutathione *S*-transferase (GST) is a family of enzymes and compress approximately a 4% of the total component of the ESPs (11). Their primary function is the detoxification of electrophiles and xenobiotics, which is essential for the parasite’s survival. In a previous study, we demonstrated that a single intraperitoneal injection with a purified native *F. hepatica* GST (nFhGST) mainly comprised of mu-class GST isoforms, administered 30 min prior to a lethal dose of LPS, significantly suppressed the cytokine storm and prevented the mortality rate either in C57BL/6 or BALB/c septic-mice (1). Knowing that the peritoneal cavity of mice is rich in macrophages, which exhibit a relevant role in the defense against microbial pathogens, tissue repairment and maintenance of homeostasis (12), we hypothesized that nFhGST could suppress the inflammatory responses by modulating the dynamic of these cells.

The macrophage activity is determined by unique signals that trigger their differentiation into distinctive phenotypes. These phenotypes differ in terms of receptor expression, effector functions and cytokine secretion. Macrophages can be classified into classically activated macrophages or M1-type macrophages and alternative activated macrophages or M2-type macrophages (13, 14). The M1-type phenotype develops in response to pro-inflammatory cytokines such as IFN-γ, and microbial or bacterial products such as LPS (13, 14). This phenotype is associated with a Th1-immune response; therefore, this phenotype induces the production of pro-inflammatory cytokines such as TNF-α. In contrast, the M2-type develops in response to an anti-inflammatory immune response, for example, in infections with parasitic helminths, and is activated by an interleukin-4/IL-13 dependent signaling pathway (13, 14). Recruitment of alternative activated macrophages (M2 type macrophages) has been observed in a various helminth infection, including *F. hepatica* (3, 5, 15–17). The present study aimed to investigate the impact of nFhGST on the dynamic of peritoneal macrophages *in vivo* and the activation and function of macrophages *in vitro*.

## Results and Discussion

### nFhGST suppresses the LPS-induced production of IL-10 and pro-inflammatory cytokines while altering the dynamic of large peritoneal macrophages in septic mice

In a previous study, we demonstrated that a single injection of 200μg nFhGST administered intraperitoneally to mice of different genetic backgrounds prior to exposure to a lethal dose of LPS was enough to prevent the cytokine storm and significantly increase the survival rate in animals exposed to lethal dose of LPS (1). In the present study, we replicated these results using BALB/c mice and investigated the effect of nFhGST on the population of peritoneal macrophages. As our results show, animals that only received the LPS injection had significantly higher levels of pro-inflammatory cytokines such as IL-12 (*p*=0.0008), IFN-γ (*p*<0.0130) and TNF-α (*p*=0.0011) than those observed in the PBS-injected control group or those only injected with nFhGST (IL-12 *p*=0.0005, IFN-γ *p*=0.013, TNF-α *p*=0.0006). IL-12 is naturally produced by dendritic cells and macrophages and is involved in the differentiation of naïve T-cells into Th1 cells (18). IFN-γ is predominantly produced by NK-cells during the innate immunity and by CD4 and CD8 T-cells during the adaptive phase of immune responses, and it has been persistently detected in persons who died of sepsis (19). TNF-α cytokine is considered an endogenous pyrogen, and its dysregulation and detection in plasma or serum of patients or animal models is a predictor of sepsis/septic shock (19). Although nFhGST does not induce any pro-inflammatory cytokines, it significantly suppressed all these cytokines (IL-12 *p*=0.0017, IFN-γ *p*=0.0144) and TNF-α *p*=0.0025) in animals that received the LPS-challenge (**Fig. 1**). It was interesting to note that nFhGST does not induce IL-10, which is a cytokine with immune-regulatory properties that was initially considered a Th2 cytokine. Conversely, animals that received the LPS-injection showed enhanced levels of IL-10 in serum, which were significantly different from those observed in the PBS-control animals (*p*=0.0003) or those injected only with nFhGST (*p*=0.0009). It is expected that during sepsis/septic shock, secretion of IL-10 could limit and terminate the inflammatory responses (20). However, recent studies have demonstrated that IL-10 can be secreted by CD4+ T regulatory cells (Tr1 cells) or Th1 cells (20) and that, at high levels could induce a state of immunoparalysis leading to an uncontrolled infection (21). IL-10 is currently considered a key cytokine in the pathophysiology of sepsis (2) and enhanced levels of IL-10 in serum have been associated with severe sepsis and a main risk factor for fatal outcomes during the condition (22) and indicate that individuals are experiencing a profound immunosuppression. The observation that nFhGST was able to significantly suppress the levels of LPS-induced IL-10 (*p*=0.0020) (**Fig. 1**) is highly relevant and suggests that nFhGST can exert a dual role of minimizing the progression of sepsis pathogenesis and preventing the immunosuppression state in the host. In agreement with the suppression of all pro-inflammatory cytokines, the abdominal cavity of animals that received nFhGST + LPS showed a gross macroscopic healthy appearance that was very similar to those observed within animals injected with PBS or nFhGST only (**Fig. 2**).

**Figure-1.**
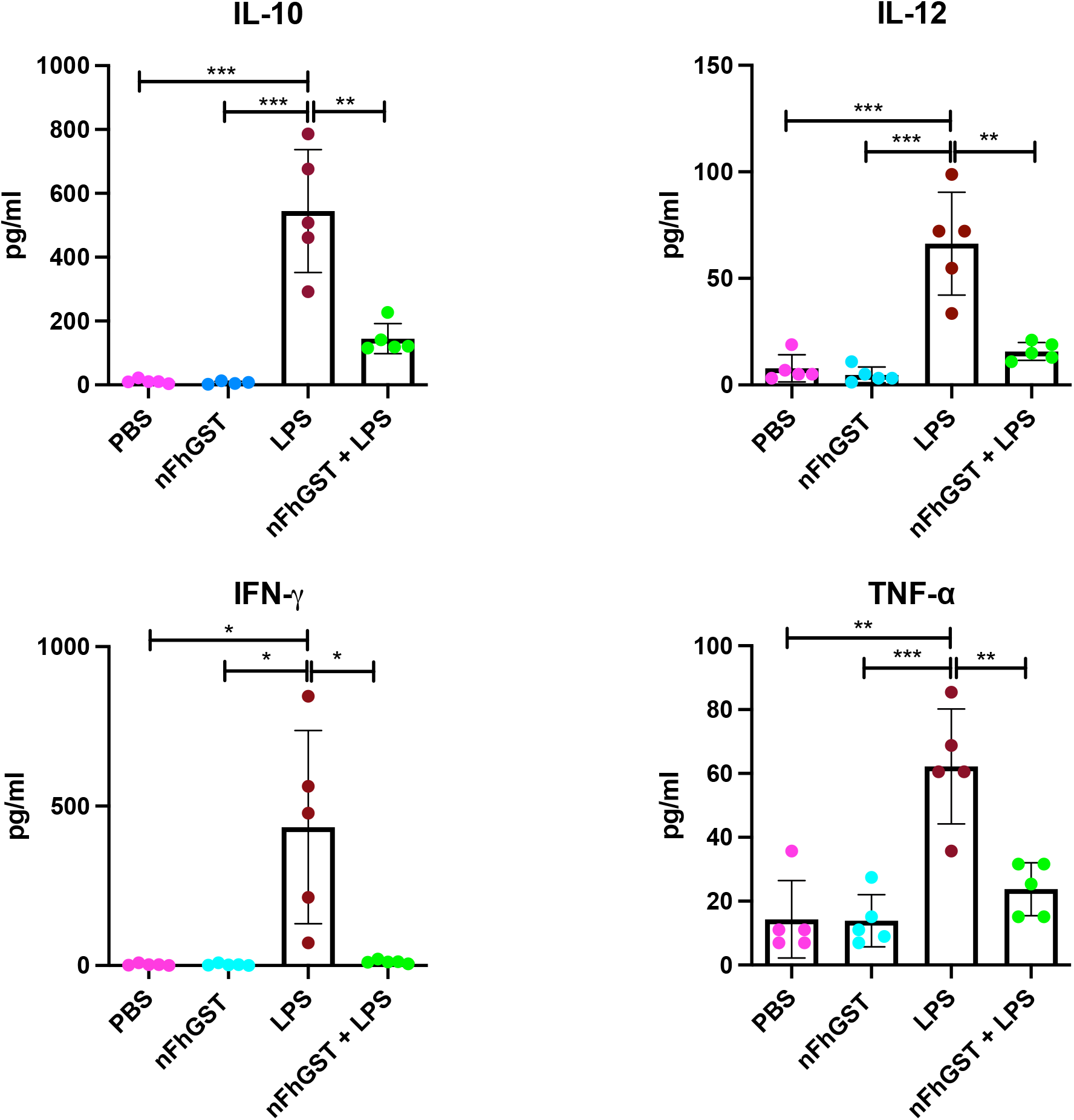
nFhGST suppresses inflammatory cytokines *in vivo* in a murine model of sepsis. BALB/c mice were allotted into groups of 5 (n=5) animals each and injected i.p. with 200μg nFhGST, 7mg/kg body wt. LPS, or 200μg nFhGST + 7mg/kg body wt. LPS. Animals injected with PBS were used as negative controls. Blood samples were taken by orbital bleeding 18 after LPS injection. Results show that nFhGST significantly reduced the LPS-induced levels of IL-10 (\*\**p*<0.0020), IL-12 (\*\**p*<0.0017), IFN-γ (\**p*<0.0144), and TNFα (\*\**p*<0.0025). Data shown herein correspond to serum samples tested in duplicate.

**Figure-2.**
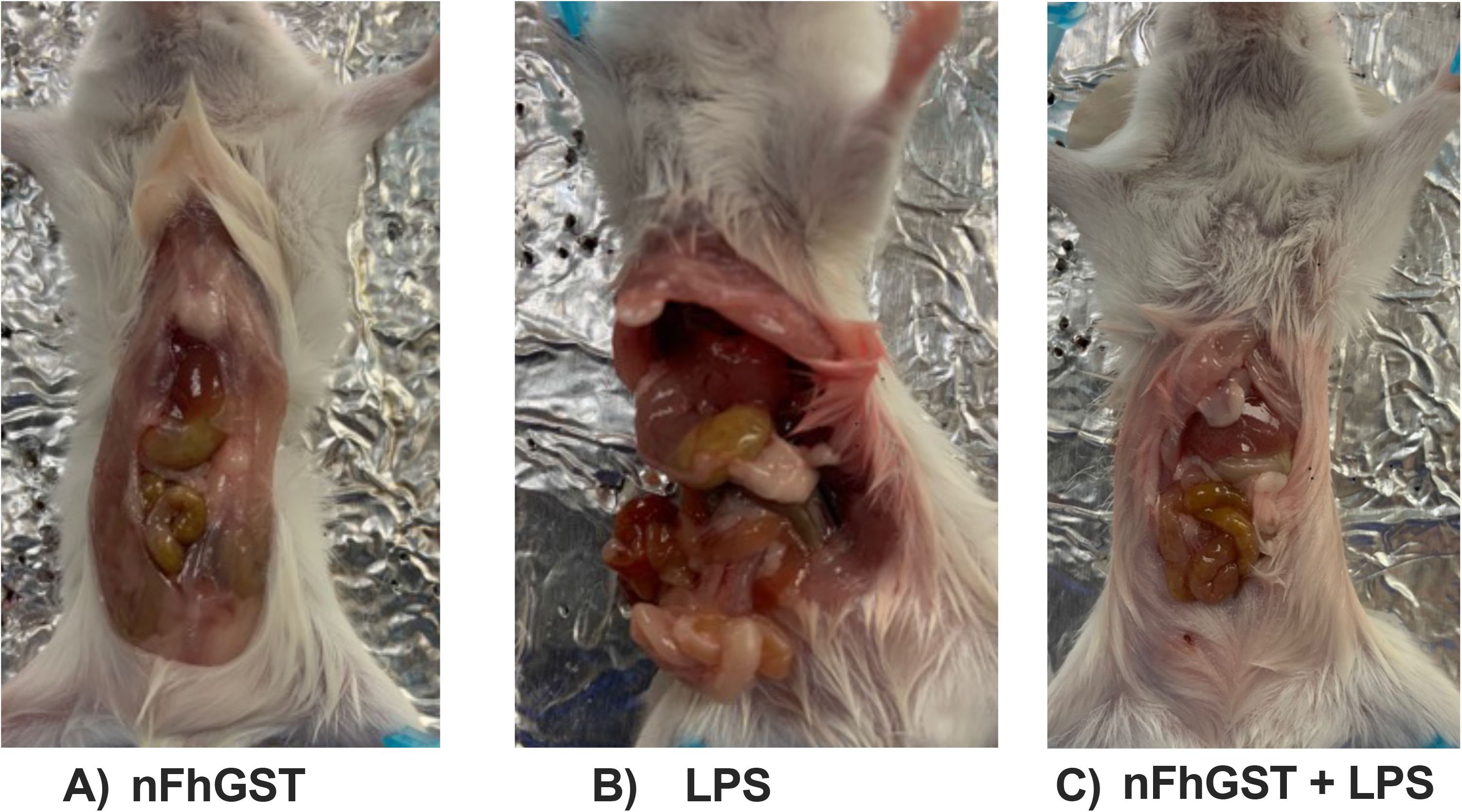
Macroscopic assessment of peritoneal cavity. BALB/c mice injected (i.p) with **(A)** nFhGST (200μg), **(B)** LPS (*E. coli* O111:B4 [7 mg/kg]), or **(C)** nFhGST 1 hour before LPS injection. Animals were sacrificed 18 hours after the LPS injection. Pictures show that the peritoneal cavity of animals treated with nFhGST or nFhGST + LPS shows a healthy appearance compared to those only injected with LPS.

Knowing that peritoneal macrophages play important roles in the tissue homeostasis and contribute to the inflammation during pathogenic infection (12), we wanted to ascertain the effect that nFhGST exerted on the abundance of these cells in the PerC. There are two different types of macrophages in the PerC of mice, which classify according to their size as small peritoneal macrophages (SPMs) and large peritoneal macrophages (LPMs) (23). LPMs are characterized by expressing high levels of F4/80 and CD11b markers and low levels of major histocompatibility complex class II (MHC-II). SPMs express lower levels of F4/80 and CD11b markers and high levels of MHC-II (23). Under steady state, LPMs and SPMs represent the 30-35% of the total peritoneal cells, whereas B1 cells constitute most of peritoneal cells along with NK-cells, lymphocytes, and B2 cells (24). It has been demonstrated that after occurring an infection or inflammatory stimuli only macrophages change in number and frequency in the PerC (23, 25). Immediately after the inflammatory stimulus, the number of LPMs reduces drastically in the PerC, a phenomenon that has been called ‘the macrophage disappearance reaction’. The disappearance of LPMs is accompanied by an increase in the number of SPMs, which it is believed increase their number due to the influx of monocytes of blood towards the peritoneum. SPMs have as their main function to secrete pro-inflammatory mediators (23, 26) and thereby are considered inflammatory macrophages (M1-type) (12). It is assumed that the disappearance of LPMs from the PerC after the inflammatory stimulus is caused by the migration of these cells to the omentum, where they proliferate and mature. After a short time, LPMs return to the PerC to control the inflammatory process (12, 23, 26). To investigate whether nFhGST could exert any influence on the dynamic of LPMs and SPMs, we washed with PBS the PerC of mice injected with PBS, LPS, nFhGST, or LPS after the nFhGST treatment to collect and labeled cells with specific anti-CD11b and F4/80 antibodies excluding from analysis dead cells, lymphocytes, B1 and B2 cells and NK cells. We found that mice that only received the LPS injection had the fewest number of LPMs, which is consistent with the macrophage disappearance reaction phenomenon. Animals that were injected with PBS had 5.78-fold more LPMs than the LPS-injected animals, and animals that were injected with nFhGST had 12.9-fold more LPMs than the LPS-control group. Importantly, LPS-injected animals that received the treatment with nFhGST had 4.87-fold more LPMs than the LPS-control group (**Fig. 3**), and these differences were found significant (*p*=0.0003), which indicates that nFhGST not only prevent the ‘disappearance’ of macrophages from PerC but becomes these cells the most abundant subpopulation in the PerC of septic animals. However, when we analyzed the SPMs subpopulation, we were not observing any change regarding the relative number of SPMs among experimental groups. Knowing that LPMs assume a role in the maintenance of PerC physiological conditions by adopting an anti-inflammatory phenotype (M2-type) (12) and that the nFhGST-treated animals with an increased number of LPMs showed significant suppression of all pro-inflammatory cytokines, it is possible to suggest that nFhGST suppress the pro-inflammatory cytokines through a mechanism based on promoting the production of LPMs to perpetuate the steady-state and homeostasis in the PerC of animals as alternative macrophages. We found a similar effect on the dynamic of LPMs in septic mice treated with Fh15, a recombinant variant of *F. he*patica fatty acid binding protein (27). This is consistent with the notion that from the beginning of *F. hepatica* infection, the parasite secretes a variety of molecules for redundantly immunomodulating the host by the recruitment of macrophages, which are alternatively activated (28).

**Figure-3.**
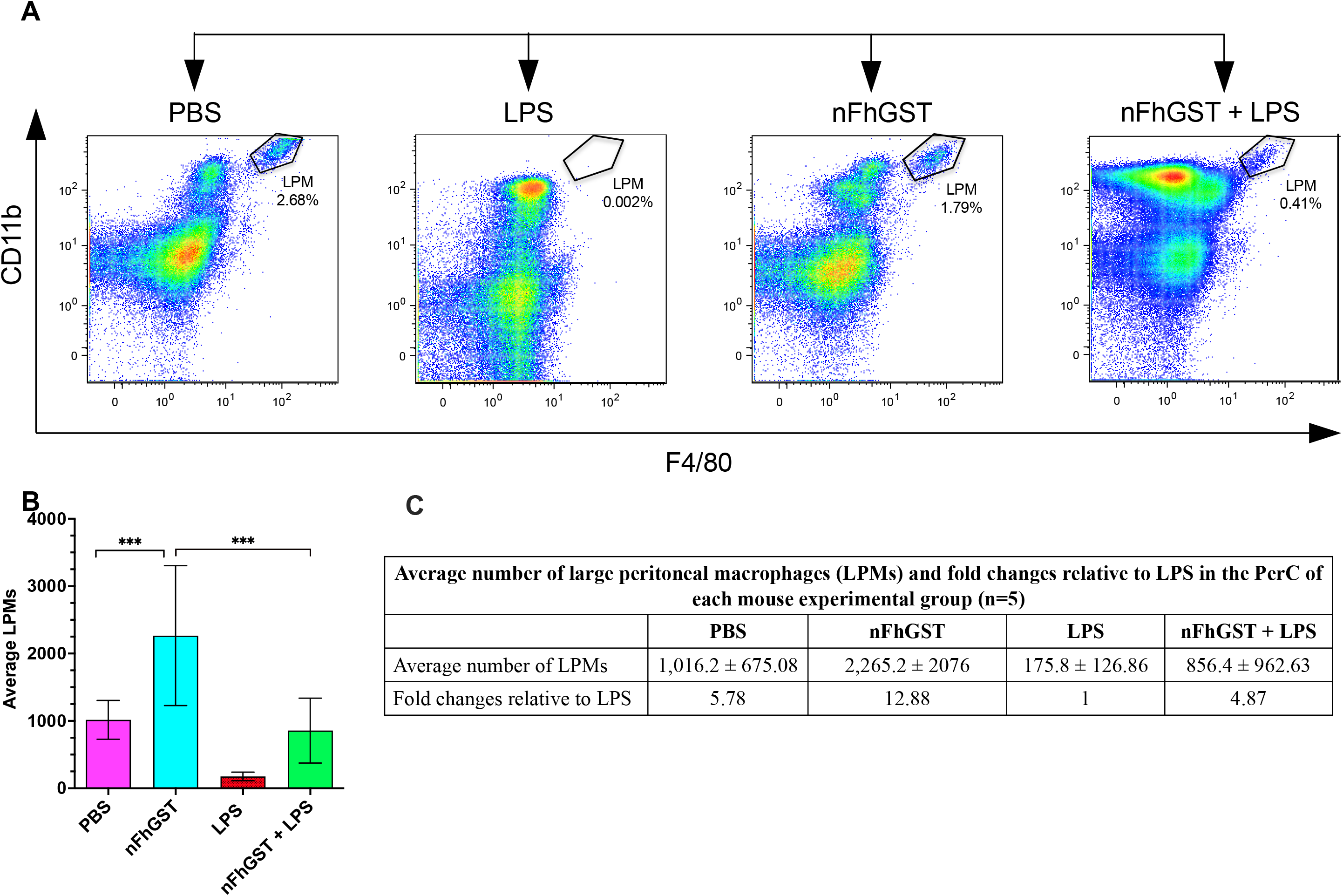
nFhGST promotes the persistence of large peritoneal macrophages (LPMs) in the peritoneal cavity in mice treated with a lethal dose of LPS. BALB/c mice were allotted into groups of 5 animals each. On the day of the procedure animals received a single intraperitoneal (i.p) injection with 200μg of nFhGST one hour before a lethal i.p. injection with 7mg /kg of body wt. of LPS (*E.coli 0111:B14*). Peritoneal macrophages (PECs) were isolated, labeled with antibodies, and analyzed by Flow cytometry. Results represent individual PECs numbers. **(A)** Flow cytometry gating representation. Cells were gated on CD11b-F4/80 cells, and LPM were identified as CD11b^+^-F4/80^+^. **(B)** Average numbers of LPMs per treatment. Bartlett’s test analysis found statistical differences between the number of LPMs from the PBS control group and LPS-injected group as well as between the number of LPMs from nFhGST and nFhGST + LPS compared to the LPS-group (\*\*\**p*=0.0003). **(C)** Fold changes of the average number of LPMs per experimental group relative to the LPS-group.

### nFhGST activates macrophages by an alternative pathway

To investigate the effect that nFhGST exerts on la activation and function of macrophages we used RAW264.7 cells, which macrophages-like cells originated from male BALB/c mice established from a tumor-induced Abelson murine leukemia virus (29). The macrophage-like cells were treated in triplicate with 10μg/ml nFhGST, 1μg/ml LPS, 20ng/ml IL-4, or 10μg/ml nFhGST for 30 min, followed by stimulation with 1μg/ml LPS and then incubated overnight at 37°C, 5% CO_2_. Cells treated with PBS were used as negative controls. As it was expected, cells stimulated with LPS, which is a strong inducer of classical activation of macrophages (M1-type) showed significantly high levels of inducible nitric oxide synthase-2 (iNOS2) and production of nitric oxide metabolites. In contrast, cells treated with IL-4, which is a strong inducer of alternative activated (M2-type) macrophages, failed in producing iNOS2 and their metabolites. iNOS2 is a key enzyme in the macrophage inflammatory response, which metabolize the arginine as a unique substrate to produce nitric oxide (NO) (30) that is potentially induced in response to inflammatory stimuli (31). Our initial analysis by confocal microscopy showed that the stimulation of macrophages-like cells with nFhGST produced low levels of iNOS2 that were not statistically different from cells stimulated with PBS or IL-4. In contrast, cells stimulated with LPS produced a significantly larger amount of iNOS2 than cells treated with nFhGST (*p*=0.0225) or IL-4 (*p*=0.0095) (**Fig. 4**). Consistent with these observations, RT-qPCR analysis also revealed significant differences (*p*= 0.0041) between the levels of iNOS2 expression in cells treated with nFhGST compared to those stimulated with LPS. The low levels of iNOS2 expression are consistent with the low levels of nitrate and nitrite detected in the supernatant of cells treated with nFhGST, which were found at the background level. However, in agreement with the high expression levels of iNOS2 induced by LPS, the levels of nitrate and nitrite detected in the supernatant of cells stimulated with LPS were significantly high (*p*=0.0002 and *p*=0.0013, respectively). Interestingly, although the treatment of cells with nFhGST prior to LPS stimulation does not exert any effect on the expression of iNOS2, it was capable of significantly reducing (*p*=0.0003) the levels of nitrate induced by LPS (**Fig. 5**).

**Figure-4.**
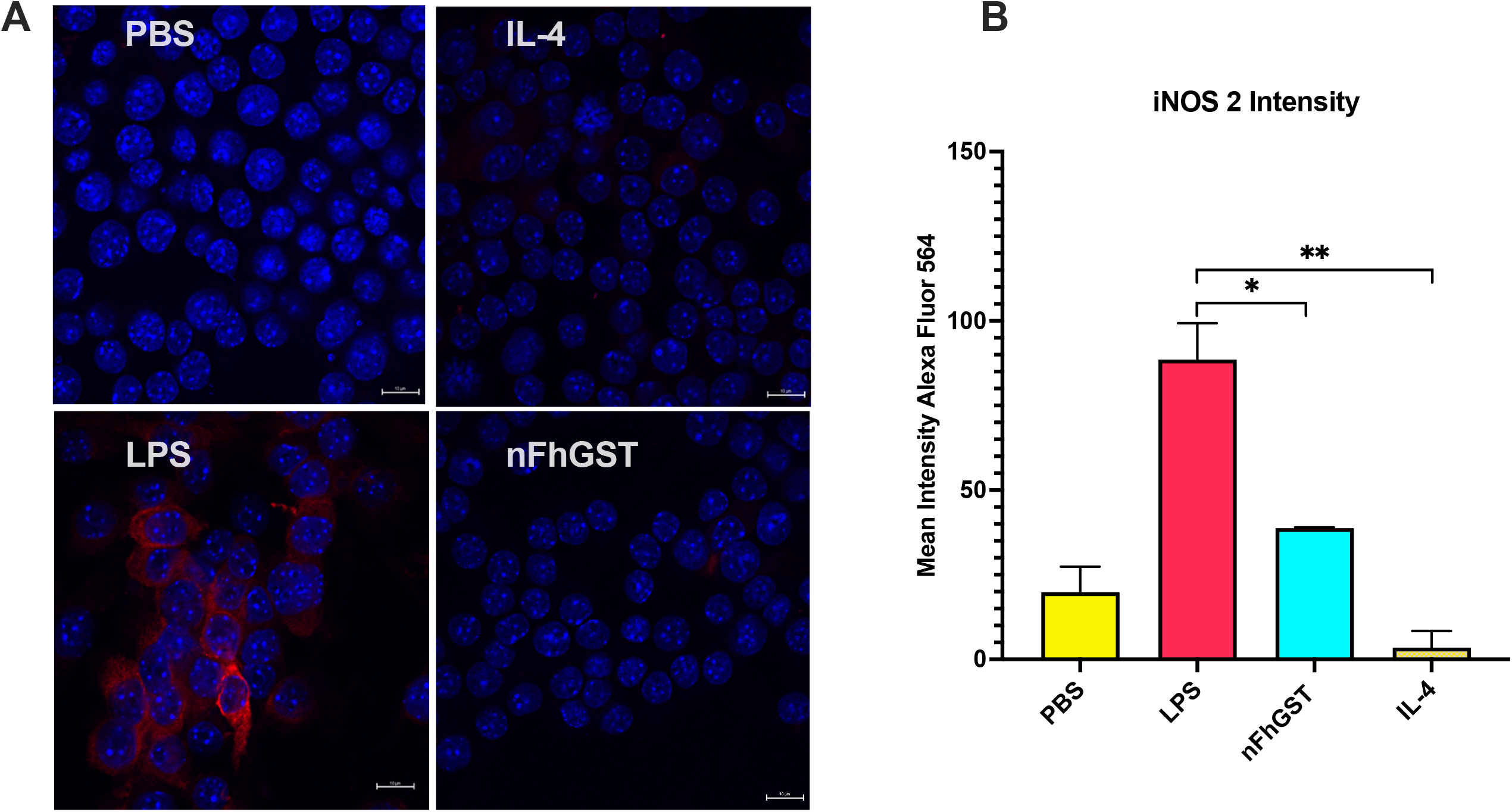
nFhGST does not produce inducible nitric oxide synthase-2. RAW264.7 cells were subcultured in 8-well glass chambers at a concentration of 30,000 cells/ well and incubated for 48 h or until 80% of confluency at 37°C, 5% CO_2_. Cells were treated overnight (O/N) with LPS (1μg/ml), IL-4 (20ng/ml), nFhGST (10μg/ml), or PBS. Cells were fixed and permeabilized and then incubated O/N at 4°C with a mouse anti-iNOS2 IgG-Alexa Fluor 564 or a mouse anti-arginase-1 IgG-FITC diluted 1:100. Cells were mounted with ProLong Diamond Antifade with DAPI (Thermo Fisher) and analyzed by confocal microscopy using Nikon Eclipse Ti Microscopy. Using the NIS-Elements Advance Research Imaging Software. **(A)** The image shows the macrophage-like cells treated with PBS, IL-4, LPS, or nFhGST where only the cells stimulated with LPS show intense red color inside cells indicating the presence of iNOS2. **(B**) The average red color intensity corresponding to Alexa Fluor 564 is significantly higher in cells stimulated with LPS compared to nFhGST (\**p*=0.0225) or IL-4 (\*\**p*=0.0095).

**Figure-5.**
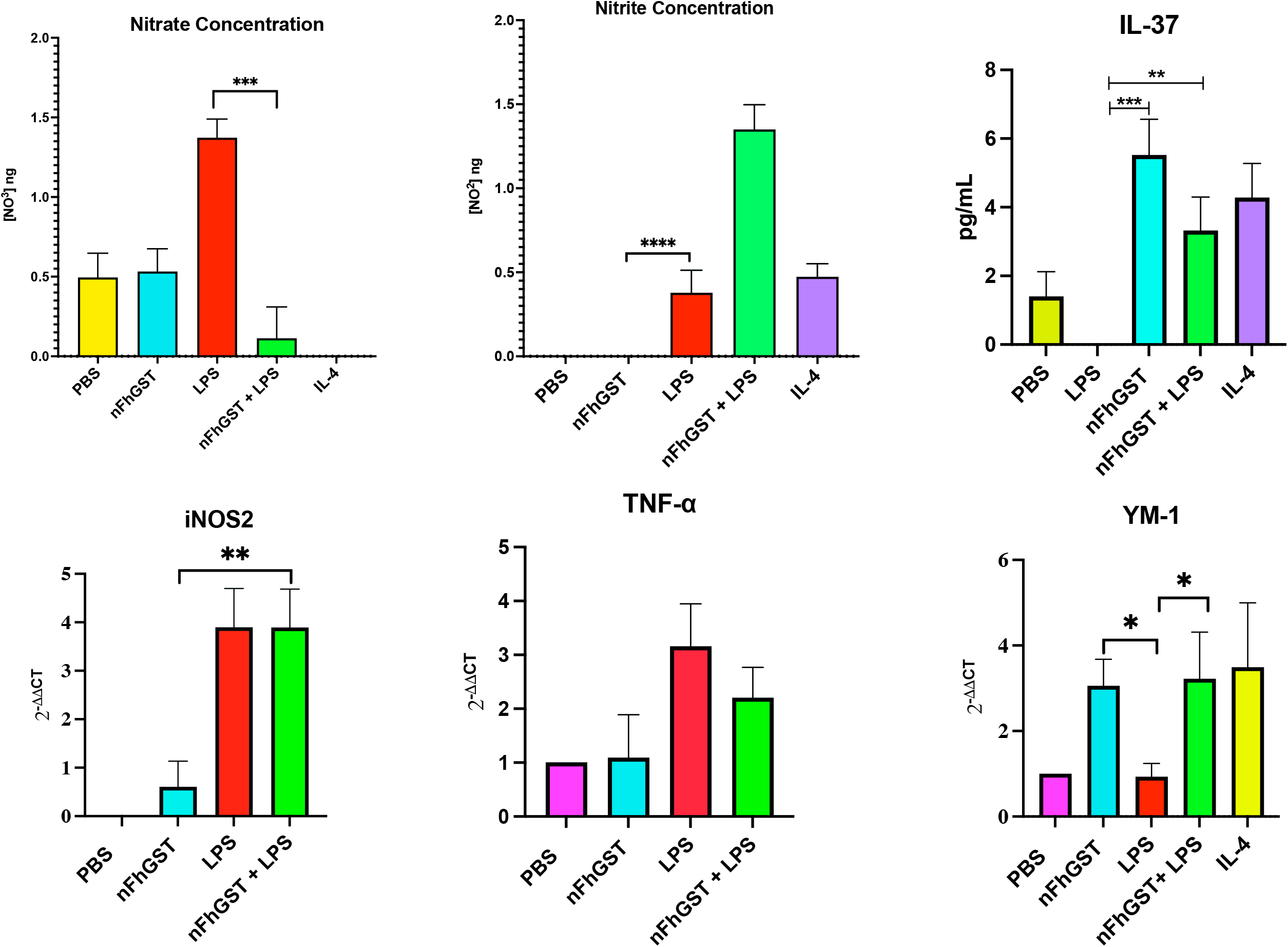
nFhGST increase Ym-1 expression and IL-37 secretion, whereas it suppresses the expression of TNF-*α*. RAW264.7 cells were cultured in a 24-well plate (4.0 x 10^5^ cells/well) for 24h and treated with LPS (1 μg/ml), IL-4 (20 ng/ml), or nFhGST (10 μg/ml). After overnight incubation at 37°C, 5% CO_2_, supernatant of cells was collected and used for quantitating levels of nitrite and nitrate using a Nitrite/Nitrate Assay or measuring levels of IL-37 ELISA following manufacturer’s instructions. Cells were detached for RNA extraction and subjected to amplification by RT-qPCR using specific primers for iNOS2, TNF-α, and Ym-1. The production of nitrate and nitrite was significantly higher in the supernatant of cells stimulated with LPS (\*\*\**p*=0.0002, \*\*\**p*=0.0013). nFhGST significantly suppressed the production of nitrite induced by LPS (\*\*\**p*=0.0003). Significantly higher levels of IL-37 were found in the supernatant of cells treated with nFhGST (\*\*\**p*=0.0005) or nFhGST + LPS (\*\**p*=0.0034). nFhGST induced significantly lower expression levels of iNOS2 (\*\**p*=0.0041), and significantly higher levels of Ym-1 expression than LPS (\**p*=0.0407).

The secretion of nitric oxide (nitrate/nitrite) and pro-inflammatory cytokines production are the most important functions tightly connected to M1-type polarization (32–34). Importantly, although nFhGST does not induce TNF-α production, it was able to reduce 1.25-fold the expression of TNF-α from macrophages-like cells stimulated by LPS, which indicates that nFhGST interferes with the development of classical activation of macrophages induced by LPS (**Fig. 5**). To ascertain whether nFhGST could be polarizing the macrophages toward M2 we proceed to investigate whether nFhGST induce the production of Arginase-1 (ARG-1), which is a recognized M2-type enzyme that uses arginine as substrate to produce ornithine and urea and play role in tissue repair and healing (35). We found that either nFhGST or LPS induced similar levels of ARG-1 and arginase activity (**see supplementary Fig. 1**). Therefore, we were unable to make the classical differentiation between M1 and M2 based on the expression of iNOS2 and ARG-1, respectively. Other studies have also reported difficulties in detecting M1 vs. M2 macrophages in *in vitro-*derived macrophages because ARG-1 is also upregulated in M1 macrophages (36, 37). Moreover, ARG-1 is expressed at very low levels in murine peritoneal cells, which makes unreliable its detection by flow cytometry (38, 39). We then proceed to determine whether nFhGST could be inducing the expression of Ym-1, which is a rodent-specific chitinase-like protein recognized as an M2-type marker involved in the modulation of macrophage activation in mice (40), the expression of Th2 cytokines, the chemotaxis of neutrophils, and other inflammatory responses (41). Ym-1 has been found overexpressed during helminth infections, including *F. hepatica*, which is recognized as the prototype of the organism’s inducer of Th2-type-immune responses with the polarization of macrophages toward M2-type (15, 42–44). As our results show, nFhGST induced high levels of Ym-1 expression that were comparable to those induced by stimulating cells with IL-4 and were significantly higher than those observed in cells stimulated with LPS (*p*=0.0407), which expressed Ym-1 at the background level. Moreover, macrophage-like cells treated with nFhGST also secreted high levels of IL-37 that were significantly higher than those produced by macrophages stimulated with LPS (*p*=0.0005) or treated with nFhGST prior to LPS stimulation (*p*=0.0034) **(Fig. 5)**. IL-37, also known as IL-1F7, is a member of the IL-1 family and has emerged as a suppressor of inflammation (45). Studies have demonstrated that IL-37 suppresses the M1 polarization of THP1-cells, which are monocytes of human origin, to inhibit inflammation (46) and that mice with transgenic expression of IL-37 were protected from LPS-induced septic shock (47). Thus, the observation that nFhGST induces the production of IL-37 from murine macrophage-like cells is highly relevant and constitutes the first report of secretion of this cytokine in murine by stimulation of a helminth-derived molecule. However, whether IL-37 induced by nFhGST could directly contribute to M2-polarization or directly contribute to switching the LPS-induced M1 phenotype to M2 is still unclear.

### nFhGST suppresses the bacterial phagocytosis by macrophages-like cells

Phagocytosis is a major function of macrophages, allowing these cells of the innate immune system to engulf and destroy foreign pathogens. To determine whether nFhGST was interfering with the phagocytic ability of macrophages-like cells, we cultured RAW264.7 cells with fluorescent-labelled *E. coli*-bioparticles in the presence and absence of nFhGST. Results demonstrated that nFhGST reduced by ∼ 45% (*p*<0.0001) the phagocytic ability of cells (**Fig. 6**). With such an inhibitory effect on macrophage function, we measured the influence of nFhGST on cell viability using an XTT assay. The results demonstrated that treatment of macrophage-like cells with PBS, LPS, or nFhGST for 24 hours did not compromise the viability of the cell (**Fig. 6**). The phagocytosis is not a function exclusive of macrophages classically activated. Several studies report that under determined circumstances, M2 macrophages can exhibit higher phagocytosis capacity than M1 macrophages and produce extracellular matrix components and regulatory cytokines like IL-10 (48, 49). However, the macrophage functions are not restricted to phagocytizing and destroying unicellular pathogens but also to regulating the immune responses (50). Macrophages are the innate immune cells that exhibit more pronounced plasticity, and currently, several M2 subtypes have been described (51), which have the role of mitigating the inflammatory responses, and promoting wound healing (52). Thus, if we consider that nFhGST promote the overexpression of Ym-1 whereas suppressing the LPS-induced TNF-α production, which denote a strong anti-inflammatory function, the theory that nFhGST induces an intermediate M2-subtype, which is opposite to the M1-type phenotype could be plausible.

**Figure-6.**
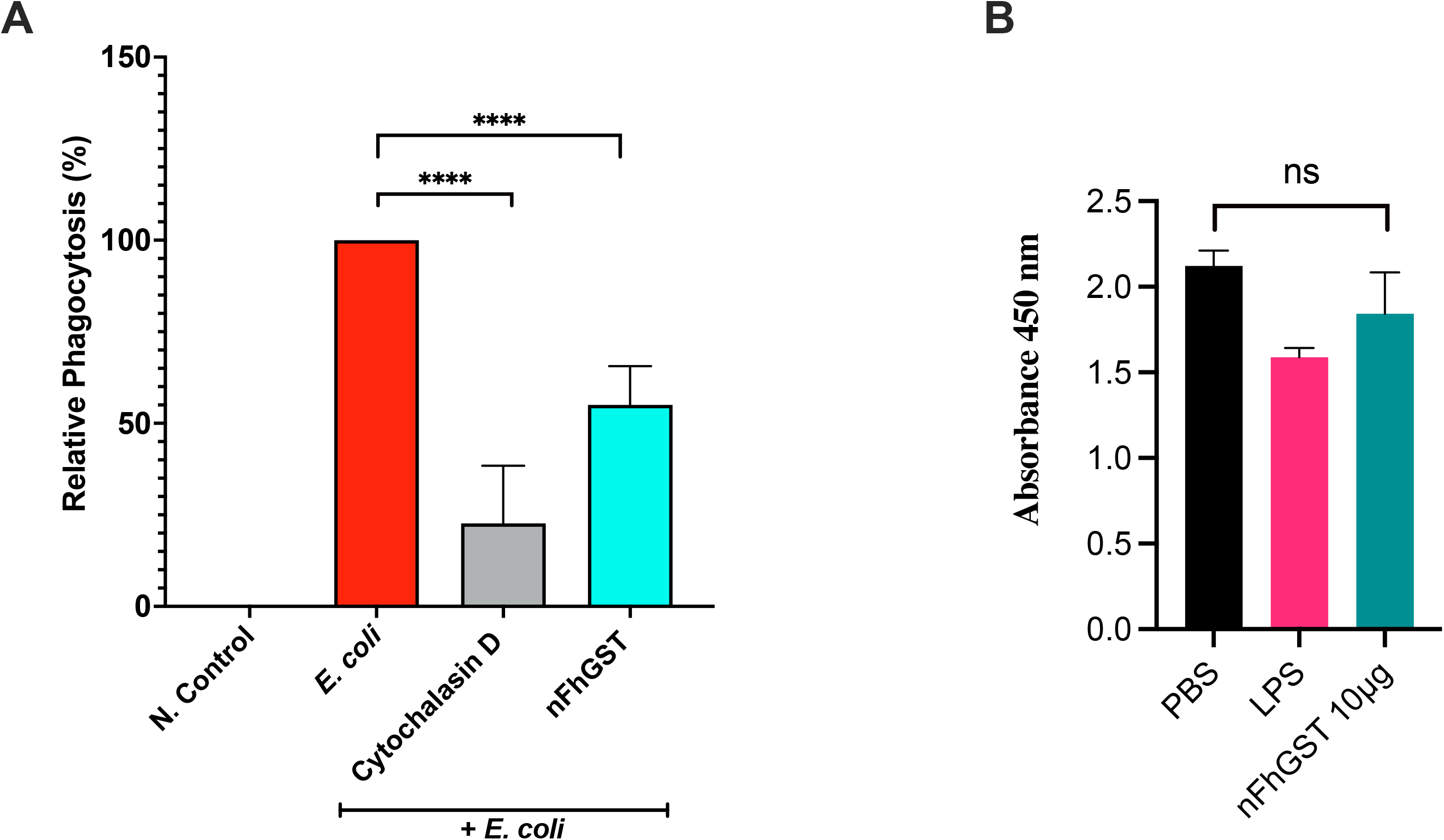
nFhGST limits the bacterial phagocytosis of macrophage-like cells. RAW 264.7 were plated at 1.0 x 10^5^ cells/well in a 96 well plate and treated for 30 min with 10μg of nFhGST, or 2μM of Cytochalasin D. **(A)** Phagocytic activity of cells was determined by adding to cells *E. coli* (K-12 strain) labeled with fluorescent dye by using a Vybrant Phagositosis assay kit. nFhGST suppressed by ∼45% the phagocytic activity of macrophages (\*\*\*\**p*<0.0005). The data shown represent the mean plus SD of four different experiments each in triplicate. **(B)** The application of an XTT-assay revealed that nFhGST or LPS treatments added to cells do not produce toxicity or interfere with the metabolism of the cells.

In summary, this study reinforces our previous observations that native *F. hep*atica glutathione S-transferase mu-class (nFhGST) exhibits powerful anti-inflammatory properties *in vivo* using a mouse model of septic shock. As part of its mechanism of action, nFhGST suppresses the cytokine storm, and although nFhGST does not induce IL-10, it can significantly suppress the LPS-induced levels of IL-10, thus mitigating the establishment of an immunosuppressive status in the animals. Moreover, nFhGST limits the phagocytic activity of cells, and suppresses the production of TNF-*α* induced by LPS *in vitro* whereas concurrently overexpress Ym-1, which is a cell marker compatible with M2-type and secrete IL-37 that is recognized as an M1-type suppressor (46).

Further studies are needed to characterize exactly the M2-type subtype that adopts the macrophages in the presence of nFhGST *in vivo* and the contribution of IL-37 to this phenotype.

## Methods

### Ethics Statement

All the animal studies were performed at the Animal Resources Center of the Medical Science Campus, University of Puerto Rico, in accordance with guidelines and protocols approved by the Ethics Institutional Animal Care and Use Committee (MSC-IACUC, Protocol No. 7870219). Female BALB/c, 6-8 weeks of age, were obtained from Charles River Laboratories. All procedures applied to the animals (bleedings, injections, and euthanasia) were performed under deep anesthesia using a cocktail of 100mg/kg body weight of ketamine + 10mg/kg of xylazine given intraperitoneally. Deep anesthesia was verified by la ack of withdrawal reflex.

### Purification of native F. hepatica Glutathione S-transferase (nFhGST)

nFhGST was purified by affinity chromatography from a *Fasciola hepatica* adult fluke extract using a 5/5 GSTrap HP column (GE Healthcare) following a pre-established protocol^9^. nFhGST is recovered in active form with a purity level of ∼96.17%, and it is comprised mostly by GST isoforms from mu-class, which was confirmed by nano-LC-MS/MS. Endotoxins were removed by using GenScript ToxinEraser^TM^ polymyxin-B (PMB) resin according to the manufacturer’s instructions, and the absence of endotoxin was confirmed using the Chromogenic *Limulus* Amebocyte Lysate QCL-1000 Assay (Lonza, Walkersville, MD). nFhGST was concentrated by AMICON Ultra-15 Centrifugal filter unit and adjusted to 1 mg/ml as determined using the Pierce protein bicinchoninic acid (BCA) method (Pierce, Cambridge, NJ). Purified nFhGST endotoxin free was stored in aliquots at −20°C until use.

### Mouse model of endotoxemia

BALB/c mice were allotted into four groups of 5 animals each and were kept under a specific pathogen-free environment (22°C, 50-60% humidity and 12h light/dark cycle) and fed with standard pellet diet and drinking water ad libitum. On the day of the procedure, animals were anesthetized to collect a small volume of blood (∼200µl) from the orbital vein using a capillary microtube to establish baseline cytokine determinations. After recovering from anesthesia, one group of animals received a single intraperitoneal (i.p) injection with 200µg of nFhGST one hour before receiving an injection of 7mg/kg of body wt. of LPS (*E. coli* 0111:B4), which is considered lethal for BALB/c mice (53). Control groups received a single injection (i.p) with PBS, nFhGST, or LPS only. Eighteen hours after the last injection, animals were anesthetized, bled out from the orbital vein, euthanized by cervical dislocation, and necropsied for collection of peritoneal exudate cells.

### Isolation of peritoneal exudate cells (PECs)

PECs were harvested from mice by washing the peritoneal cavity (PerC) with 10-20 ml of cold PBS as described elsewhere (54). PECs were centrifuged for 5 minutes at 2,200 rpm. The supernatant was discarded, and pellet cells were resuspended in 1mL of PBS, counted and adjusted to a concentration of 1.0x 10^6^ cells/ml.

### Flow Cytometry analysis of peritoneal exclude cells (PECs)

PECs isolated from experimental animals were stained and incubated for 30 min with Zombie Violet Fixable Viability Dye (BioLegend) to assess dead cells. After incubation, cells were washed with PBS, and the Fc receptor of the cells was blocked using an anti-mouse CD16/32 antibody (Fcγ R III/II, Ly-17; BioLegend). After 30 min of incubation at 4°C cells were washed with FACS Buffer (PBS containing 1% FBS) and subsequently centrifuged at 1,200 rpm for 5 min. After removing the supernatant, cells were resuspended in 100μL of antibody master mix and incubated for 30 min at 4°C. The following antibodies were used in this experiment: F4/80 (PerCP-Cy5.5 BioLegend), CD11b (APC-Cy7, BioLegend). Untreated cells were used as a non-specific staining control. Unstained cells were used to normalize the data. To separate auto-fluorescing cells from cell expression low levels of the different markers, we upper thresholds for autofluorescence by staining samples fluorescence-minus-one (FMOs) stain sets, in which the reagent of interest is omitted. Dead cells were excluded, and macrophages were identified as F4/80^+^ CD11b^+^. The acquisition was performed using a Miltenyi MACSQuant Analyzer 10 instrument, and the data was analyzed using FlowJo software (FlowJo, LLC).

### Cytokine Measurement

Serum samples collected from mice were tested in duplicate in a multiplex immunoassay using magnetic beads (Mouse Th1-Th2 8-plex Bio-Rad, Hercules, CA) using the Luminex MAGPIX® system). Data were analyzed with Bio-Plex Manager 6.1 software (Bio-Rad, Hercules, CA) using a 5-parameter logistic curve.

### Macrophage cell line passage and stimulation

RAW264.7 cells, a mouse monocyte/macrophage cell line established from a tumor in a male mouse induced with the Abelson leukemia virus, were purchased from ATCC and cultured on a T25 cell culture flask in Dulbecco’s modified minimal essential medium (DMEM) supplemented with 10% (v/v) of heat-inactivated fetal bovine serum (iFBS), 100 U/mL penicillin and 100µg/ml of streptomycin at 37°C, 5% CO_2_. The cells were passage every 2 days, and when reached 60% confluency were subculture and used in further experiments.

### iNOS2 and ARG-1 detection by confocal microscopy

RAW264.7 cells cultured as described above were detached from the T25 flask using a cell scrapper, counted, and subcultured in 8-well glass chambers at a concentration of 30,000 cells per well and incubated for 48 h or until 80% of confluency at 37°C, 5% CO_2_. Cells were treated with a pre-optimized concentrations of LPS (1μg/ml) to obtain M1 macrophages, or IL-4 (20ng/ml) for M2 differentiation. Cells were also treated with pre-optimized concentration of nFhGST (10μg/ml) or treated with nFhGST 30 min prior to stimulation with LPS. Cell treated with PBS served as controls. Cells were fixed with 4% of paraformaldehyde (PFA) for 15 min and permeabilized with 0.5% Triton X-100. Afterward, cells were incubated O/N at 4°C with a mouse anti-iNOS2 IgG antibody labeled with Alexa Fluor 564 (BioLegend)) or a mouse anti-ARG-1 IgG antibody labeled with FITC (Santa Cruz Biotechnology) diluted 1:100. Next, cells were mounted with ProLong Diamond Antifade with DAPI (Thermo Fisher) and analyzed by confocal microscopy using Nikon Eclipse Ti Microscope. Using the NIS-Elements Advance Research Imaging Software. The mean intensity of a minimum of three different images, each in triplicate generated each for Alexa Fluor 564-iNOS2 IgG antibody or FITC-ARD-1 antibody was measured.

### Measurement of Nitrate Nitrite levels produced by macrophage-like cells

RAW264.7 cells were subculture in a 24-well plate (4.0 x 10^5^ cells per well) for 24h and treated with LPS (1μg/ml), IL-4 (20ng/ml) or nFhGST (10μg/ml) as described above. After overnight incubation at 37°C, 5% CO_2_, supernatant of cells was collected and used for quantitating levels of nitrite and nitrate using a Nitrite/Nitrate Assay (Sigma Aldrich) following manufacturer’s instructions. Briefly, 40μl of all supernatants was added to four wells in a 96-well plate. Two wells were used to measure NO_2_^-^ detection, and the other two wells for the [NO_3_^-^ + NO_2_] detection. Nitrate reductase and Enzyme Co-factors were added as described in the kit protocol. After 2 h of incubation at room temperature, 50μl of Griess Reagent A was added to all wells and incubated for 5 min. After incubation another 50μL of Griess Reagent B was added to all wells and incubation was prolonged for 10 minutes. After the last incubation, the absorbance was measured at 540 nm. The levels of nitrite and nitrate were calculated based on corresponding standard curves according to the manufacturer’s instructions.

### Measurement of interleukin-37 (IL-37) levels secreted by macrophages-like cells

RAW264.7 cells were subcultured in a 24-well plate using a mouse IL-37 ELISA kit (My BioSource, San Diego, CA.). Briefly, 100μL of supernatants and standards were added in duplicate to a 96-wells plate pre-coated with a specific anti-IL-37 antibody and incubate for 2h at 37°C. After removing any unbound substances, a biotin-conjugated antibody specific for IL-37 was added and incubated for 1 h at 37°C. After washing, streptavidin conjugated horseradish peroxidase (HRP) was added to all wells, and the plate was incubated for 1h at 37°C. After another washing step, the substrate solution was added to all wells and incubated for 15-20 minutes at 37°C. The reaction was stopped, and absorbance was read at 450nm with the correction wavelength set at 540nm. A standard curve was generated by diluting the IL-37 control at concentrations ranging between 3.12pg/ml to 200pg/ml.

### RNA isolation and RT-qPCR analysis

Adherent RAW264.7 cultured in a 24-well plate and treated with LPS, IL-4, or nFhGST as described above were used for measuring the mRNA expression of iNOS2, Ym-1 and TNF-α using specific primers for each of these molecules **(Table-1)**. Total RNA was isolated from these samples according to the manufacturer’s instruction using AllPrep DNA/RNA/Protein Mini Kit (Qiagen). RNA was quantified using a Nanodrop-1000 spectrophotometer (Thermo-Scientific, USA) and amplified using Power SYBR Green RNA to CT 1-Step Master Mix (Applied Biosystems). The experiments were conducted in triplicate using the Quant Studio 3 PCR system (Applied Biosystems). A concentration of 100ng of RNA was used. Reaction was initiated with an initial incubation at 48°C for 30 min, and 95°C for 10 min followed by 40 cycles of 95°C for 15 s and 60°C for 1min. Primer concentration was optimized, and dissociation curves were generated for each primer set to verify the amplification of a single PCR product. The relative gene expression levels were calculated using the 2^−ΔΔCt^ method. The gene primers for qPCR are provided in table 1. GAPDH was an internal reference gene between different samples and expressed as fold change relative to the expression of the control (cells stimulated with PBS).

**Table-1.**
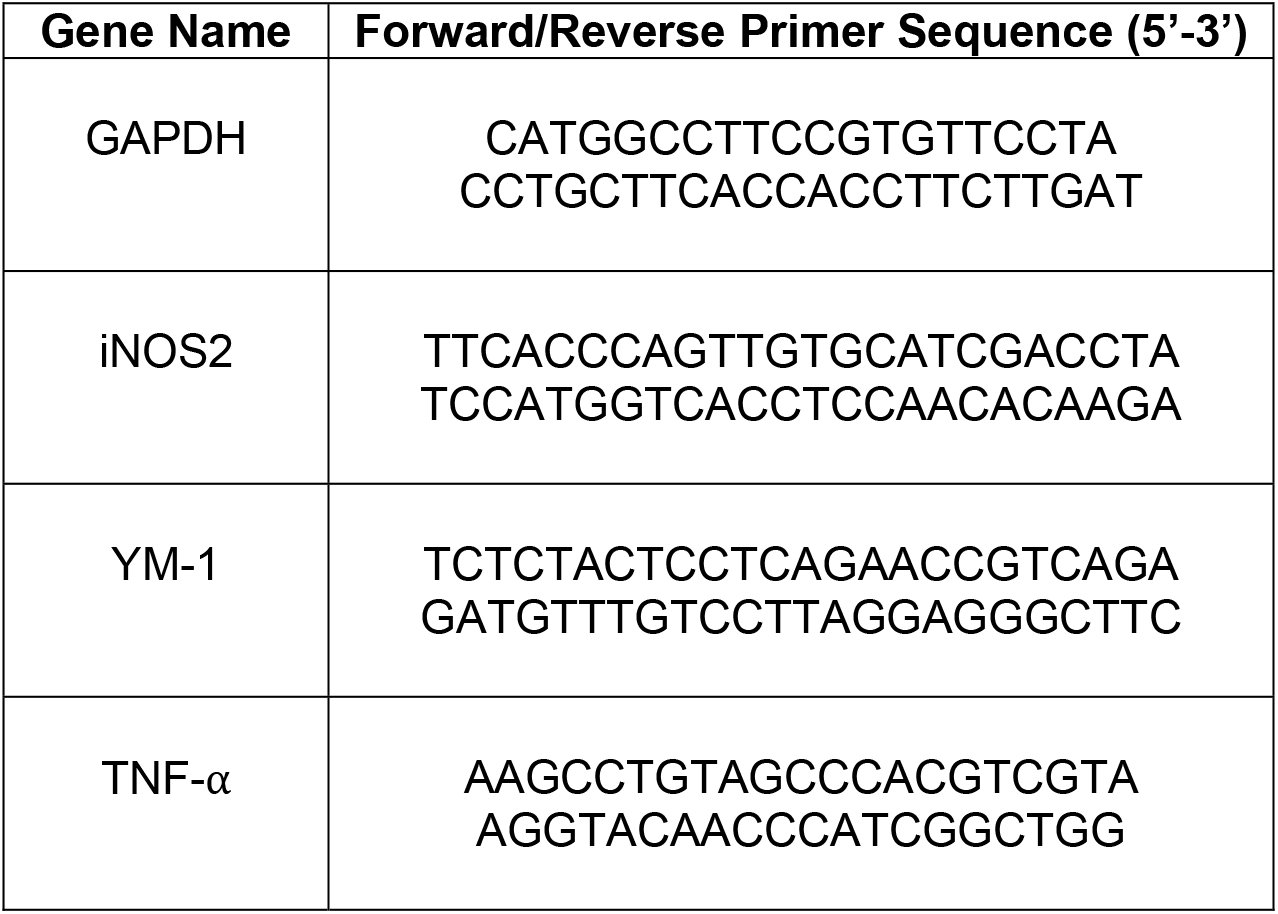
Primer sequences used in quantitative real-time polymerase chain reaction (qPCR) analysis.

### Phagocytosis Assay

The phagocytic ability of RAW 264.7 in the presence of nFhGST was measured using a Vybrant Phagocytosis Assay (Invitrogen) that uses *E. coli* (K-12 strain) labeled with fluorescein. Briefly, RAW 264.7 were plated at 1.0 x 10^5^ cells/well in a 96 well plate and treated for 30 min with 10μg of nFhGST or 2μM of Cytochalasin D. After the incubation, the treatment was removed from all wells by vacuum aspiration. Cells were then treated with 100μl of the prepared fluorescent BioParticle suspension (negative control, positive control, and experimental wells) and incubated at 5% CO_2_ 37°C for 2 hr. After incubation, BioParticle suspension was removed by vacuum aspiration, and 100μl of prepared Trypan Blue was added to all wells and incubated for 1 min at room temperature. After incubation, triptan blue was removed by vacuum aspiration. The microplate was read at 480 nm excitation and 520 nm emission. The net phagocytosis percentage and the response of the phagocytosis effector agent were calculated as described in the kit protocol calculating the Net Experimental Reading and the Net Positive Reading. The phagocytosis response to the effector was expressed as follows: % Effect= <u>Net Experimental Reading</u> x 100 Net Positive Reading. The phagocytosis response to the effector was expressed as follows:

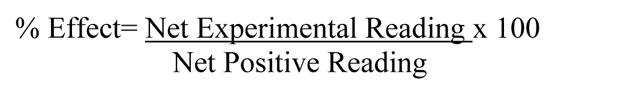

### Measurement of Cell Viability

The effect of nFhGST on the viability of RAW264.7 cells was measured by an XTT assay, according to the manufacturer’s instructions. In brief, RAW264.7 cells were cultured in a 96-well microplate at an initial density of 1 x 10^4^ cells/well for 24h. Then cells were treated with 10µg/ml nFhGST, LPS (1µg/ml) or co-treated with nFhGST + LPS for an additional 18 h under normal cell culture conditions. Equal volumes of XTT Developer Reagent and Electron Mediator Solution (Abcam ab232856) were combined to make the XTT mixture. Then the 10µl of the XTT mixture (sodium 3’-[1-phenylaminocarbonyl-)-3,4 tetrazolium]-bis(4-methoxy-6nitro) benzene sulfonic acid hydrate) were added to each well. Cells were incubated for additional 2 hours at 37°C in a CO_2_ incubator. The absorbance was determined at 450 nm using a microplate reader (Thermo Scientific Multiskan FC). All experiments were carried out in four biological replicates.

### Statistical analysis

All determinations were performed in duplicate or triplicate, and each independent experiment was replicated twice. For comparisons between two groups in all the experiments an unpaired Student’s *t*-test was used. All statistics were done with GraphPad Prism 8.0 (San Diego, CA).

## Acknowledgments

This research was supported by Grant number SC1AI155439-01 from the National Institute of Allergy and Infectious Diseases and grants G12MD007600, R25GM061838 and 5R25GM061151 from the National Institute on Minority Health and Health Disparities and by the National Institute of General Medical Sciences (NIGMS) of the National Institutes of Health (NIH) under Award Number P20GM103642. We wish to thank Lcd. Bismark Madera, Dr. Dina Bracho, and the Neuroimaging and Electrophysiology Facility (NIEF) staff for the immunofluorescence images and analysis. The content is solely the responsibility of the authors and does not necessarily represent the official views of the National Institutes of Health.

## Authors declare no conflict of interest

## Author Contributions Statement

**BV:** Designed the experiments, performed the experiments, statistical analysis, prepared figures and drafted the manuscript.

**AAR**: Collaborated in the *in vivo* experiments and revised the manuscript.

**CRJ**: Collaborated in the design and analysis of the *in vivo* experiments and revised the manuscript.

**LBM**: Provided Bioplex technology and collaborated with the cytokine analysis and revised the manuscript.

**AME:** As PI, provided the funding and laboratory space where the experiments were performed. Designed the experiments, participated in the data analysis, prepared figures, revised the manuscript, and wrote the discussion.

**Supplementary Figure-1. Macrophages-like cells express similar levels of ARG-1 when are stimulated with LPS, IL-4, or nFhGST. (A)** RAW 264.7 cells were subcultured until 80% confluency in an 8 well glass chamber (30,000 cells/ml) for 48h at 5% CO_2,_ 37°C. Cells were stimulated with LPS (1μg/ml), nFhGST (10 μg/ml), or IL-4 (20 ng/mL). After overnight incubation, cells were fixed, permeabilized, and stained with Arginase-1 FITC antibody (1:100) and visualized using a Nikon Eclipse Ti-Microscope and analyzed using NIS-Elements Advance Research Imaging Software. **(B)** The mean intensity fluorescence of the FITC-ARG-1 antibody was measured. Results represent the means ± SD from a minimum of three experiments, each in triplicate. \*\*\*\**p*<0.0001 and \*\*\**p*<0.0006.

